# Environmental impacts of cultured meat: A cradle-to-gate life cycle assessment

**DOI:** 10.1101/2023.04.21.537778

**Authors:** Derrick Risner, Yoonbin Kim, Cuong Nguyen, Justin B. Siegel, Edward S. Spang

## Abstract

Interest in animal cell-based meat (ACBM) or cultured meat as a viable environmentally conscious replacement for livestock production has been increasing, however a life cycle assessment for the current production methods of ACBM has not been conducted. Currently, ACBM products are being produced at a small scale and at an economic loss, however ACBM companies are intending to industrialize and scale-up production. This study assesses the potential environmental impact of near term ACBM production. Updated findings from recent technoeconomic assessments (TEAs) of ACBM and a life cycle assessment of Essential 8™ were utilized to perform a life cycle assessment of near-term ACBM production. A scenario analysis was conducted utilizing the metabolic requirements examined in the TEAs of ACBM and a purification factor from the Essential 8™ life cycle assessment was utilized to account for growth medium component processing. The results indicate that the environmental impact of near-term ACBM production is likely to be orders of magnitude higher than median beef production if a highly refined growth medium is utilized for ACBM production.

## Introduction

Livestock production is an integral component of the global food system, providing staple proteins (milk, eggs, and meat) consumed worldwide, contributing to crop productivity via utilization of manure as fertilizer, and providing critical nutrition and income to underprivileged households in low to middle income countries (Gilbert et al., 2018; Robinson et al., 2011). Global meat production has increased from 70.57 million tonnes in 1961 to 337.18 million tonnes in 2020, though the consumption of different meat sources is highly regionalized (FOA, 2022; Ritchie et al., 2019). In 2020, beef and buffalo meat accounted for ∼22% of global meat production, and poultry and pork accounted for ∼39% and ∼32% of worldwide meat production, respectively (FOA, 2022; Ritchie et al., 2019).

Looking forward, the overall demand for meat is expected to double by 2050 (Food and Agriculture Organization of the United Nation (FAO), 2019), and this trend has raised concerns about the environmental impact of scaling up meat production to meet these expected demands. When the top three livestock production systems are examined from an environmental perspective, beef is the most impactful per kilogram, though this value varies significantly by production system (Poore & Nemecek, 2018). The environmental impact of beef production includes greenhouse gas emissions (GHG) from enteric fermentation and manure, nutrient loading in the nitrogen and phosphorus cycles, reduction in biodiversity from overgrazing, and deforestation from land-use change (Gilbert et al., 2018; Steinfeld et al., 2006).

Multiple life cycle assessments (LCAs) have examined different beef production systems and the global warming potential or GWP (kg of carbon dioxide equivalent, CO_2_-eq) was the most highly utilized environmental metric for these assessments (de Vries et al., 2015). This impact is then normalized by the functional unit of the beef product (e.g. live weight, carcass weight and boneless meat), which varies across studies. For example, skeletal muscle is only one product produced from a slaughter facility (Desjardins et al., 2012). Approximately 78.3% mass of the animal is utilized as primal cuts of meat (37.8%), rendering products (32.8%), raw hide, (4.9%) and offal (3.2%) in the United States and Canada (Desjardins et al., 2012). A 2015 review of beef LCAs reported a range of 7.6 kg (live weight) to 29.7 kg (carcass weight) of CO_2_e per kg of beef (de Vries et al., 2015).

The reported values in the literature vary significantly due to differences in functional unit, as mentioned above, but also by the production system (e.g. origin of calf, organic vs. non-organic, and type of diet), and geographic location (de Vries et al., 2015). A study that examined the environmental impact of multiple foods at the retail level indicated GHG emissions ranged from 9.6 to 432 kg of CO_2_e for each kilogram of fat and bone-free meat and edible offal (FBFMO) produced (Poore & Nemecek, 2018). The reported GHG emissions from meat produced from a beef herd (cattle raised with primary purpose of meat production) ranged from 35-432 kg of CO_2_e per kg of FBFMO. After statistical analysis, the mean and median for the beef herd was 99.5 and 60.4 kg of CO_2_e per kg of FBFMO. The greenhouse gas emissions from FBFMO produced from dairy herds ranged from 9.6 to 73.9 kg of CO_2_e per kg of FBFMO. The mean and median of the greenhouse gases produced from FBFMO production from dairy herds was 33.4 and 34.1 kg of CO_2_e per kg of FBFMO, respectively (Poore & Nemecek, 2018). The relative closeness of the mean and median indicate fewer outliers for dairy herd produced FBFMO. Due to the potential environmental impacts of increased beef production and animal welfare concerns, beef production has been identified as a large-scale food production system that could be modified, significantly curtailed, or even eliminated (McMichael et al., 2007; Pierrehumbert & Eshel, 2015).

### Alternative Protein Products and Animal Cell-Based Meat (ACBM)

Several methods or system alternatives have been proposed to reduce the environmental impact of human-consumed proteins including alternative protein production, regenerative agriculture, and bovine methane reduction (“clean cow”) efforts (Cusworth et al., 2022; Min et al., 2022; Molfetta et al., 2022). During the last five to ten years, alternative proteins or meat alternatives have gained popularity with a multitude of stakeholders. These stakeholders have coalesced around this concept to augment or replace conventional beef production (Tziva et al., 2020). The interest of these stakeholders is multifaceted and includes concerns for animal welfare, environmental concerns, and/or profit-seeking motivations. The multifaceted nature of these stakeholders can be illustrated by non-profit groups like The Good Food Institute which exhibits interests in a mix of social activism, scientific inquiry, and financial investment.

Alternative proteins can be broadly categorized into three distinct categories: plant-based proteins, fermentation-based proteins, and animal cell-based meat (ACBM) or “cultured meat” (Asgar et al., 2010; Tziva et al., 2020). Plant-based and fermentation-based proteins are currently commercially available, and these products have been for several decades (Ex. Tofurky and Quorn^®^, respectively) (Tziva et al., 2020). The core concept of ACBM production is that animal cells such as pluripotent stem cells can be proliferated in industrial scale bioreactors (>1,000 L), differentiated into a variety of cell types (e.g. adipocytes, myotubes, fibroblasts), and then processed for human consumption in place of conventionally produced meat (Humbird, 2021; Risner et al., 2020). At the time of this writing, no ACBM products are produced at a large enough scale to be considered commercially available. The authors acknowledge the current small-scale production of ACBM products in Singapore, however these products still utilize animal serums such as fetal bovine serum and are not widely available (Hasiotis, 2022). Additional challenges related to organoleptic quality of these novel products are also evident (Fraeye et al., 2020).

Despite the highly limited availability of ACBM products, investment in ACBM companies has continued to increase with a total investment of over $2 billion at the time of writing (Turi, 2021). This investor excitement is likely linked to analyst’s reports which are bullish on meat alternatives with some reports predicting a 60-70% displacement of ground beef by 2030-2040 (Suhlmann et al., 2019; Tubb & Seba, 2019). More recent reports seem to be more modest with their predictions of replacing a half of a percent of conventional meat products with ACBM products by 2030 (Brennan et al., 2021). With 12.6 billion kg of beef produced in the United States in 2021 (Maples, 2021), even this more conservative estimate of predicted displacement would have a massive impact on the food system.

### Existing Technoeconomic Assessments of ACBM

Given the reported potential impact of ACBM production, researchers at the University of California, Davis (UC Davis) published a preliminary, peer-reviewed techno-economic assessment (TEA) of ACBM that examined the core capital and operating expenditures required to produce ACBM at scale (Risner et al. 2020). Given the uncertainty of auxiliary processes (i.e. scaffolding, product forming or shaping, etc.) the TEA focused on the core cell proliferation and differentiation processes in production scale bioreactors. The production scale bioreactors represented the system capital costs and the variable operating expenditures included ingredients, raw materials, some utilities, and labor cost. The Risner et al. TEA included Essential 8™ (E8) as the animal cell growth medium for their model. E8 is a defined growth medium designed for stem cell research and had been previously suggested as a growth medium which could be scaled and slightly modified for industrial production of ACBM (Chen et al., 2011; E. A. Specht et al., 2018; L. Specht, 2019). The authors believe that use of E8 or similar refined growth medium will be necessary given *in vitro* animal cells sensitivity to media impurities in comparison to yeast or bacterial cells.

Given the uncertainty inherent to modeling an emerging technology, the Risner et al. TEA included an assessment of four potential scenarios for the production of 122 million kg of ACBM (wet cells) or alternatively, 36.6 million kg of dry cells and 25.62 million kg of protein. Scenarios 1 and 4 represented “bookend” scenarios where Scenario 1 represented the initial state of ACBM production mirroring the economics of early proof of concept demonstrations and Scenario 4 represented achieving the physical and biological limits of the bioreactor (thus, not an operationally realistic scenario for actual ACBM production). Scenarios two and three represented “midpoint” scenarios where a few particularly critical cost hurdles were overcome.

Shortly after the Risner et al. TEA was published, a more complete TEA commissioned by Open Philanthropy was peer reviewed and published in *Biotechnology and Bioengineering* (Humbird, 2021). This TEA examined a complete production system and included all the equipment that would be necessary at a scale of 100 million kg of ACBM produced per year. The Humbird TEA examined a more simplified growth medium with commodity level pricing and refinement for the carbon source. The Humbird TEA also utilized chemical engineering scaling equations to estimate costs at scale.

### The Endotoxin Challenge

These TEAs highlighted many of the technical challenges related to ACBM production, but growth medium refinement was identified as one of the most important considerations for nearterm analysis. One aspect of this refinement is the endotoxin reduction/removal for each growth medium component. Endotoxins, also known as lipopolysaccharides (LPS) are a critical component of the outer membrane of Gram-negative bacteria. Endotoxins contain a hydrophilic polysaccharide fraction, which is covalently bonded to a hydrophobic lipid known as lipid A (Magalhães et al., 2007). Gram negative bacteria are ubiquitous to the environment and are commonly found in tap water (Vaz-Moreira et al., 2017). In cell culture the presence of endotoxin can have a wide variety of effects. For example, at an endotoxin concentration as low as 1 ng/ml it reduced pregnancy success rates by 3 to 4-fold during *in vitro* fertilization of human IVF embryos (Dawson, 1998; Fishel et al., 1988; Snyman & van der Merwe, 1986). Gram negative bacteria shed small amounts of endotoxin into the environment when they proliferate and shed large amounts when they are inactivated (Corning, 2020).

Animal cell culture is traditionally done with growth medium components which have been refined to remove/reduce endotoxin (Corning, 2020). The method of endotoxin reduction or elimination is highly dependent upon the properties of the substance being purified (EMD Millipore, 2012). There are a multitude of methods employed for the separation of endotoxin from growth medium components and these include use of LPS affinity resins, two-phase extractions, ultrafiltration, hydrophobic interaction chromatography, ion exchange chromatography, and membrane adsorbers (Magalhães et al., 2007). In turn, the use of these refinement methods contributes significantly to the economic and environmental costs associated with pharmaceutical products since they are both energy and resource intensive (Wernet et al., 2010).

### The Limitations of existing ACBM LCAs

Previously conducted LCAs of ACBM have significant limitations, namely, the high levels of uncertainty in their results and the lack of accounting for endotoxin removal. Despite study authors clearly reporting high levels of uncertainty in their LCAs, the results are often cited as clear evidence for the sustainability of ACBM production. The potential environmental impact of producing ACBM has been evaluated to be less than conventionally produced beef in previously conducted LCAs of ACBM (Mattick et al., 2015; Tuomisto et al., 2014; Tuomisto & Teixeira de Mattos, 2011). An often-cited LCA of ACBM production claims 1.9-2.2 CO_2_eq GHG emissions are emitted and 26–33 MJ energy will be utilized per kg of ACBM produced. This assessment is based on utilizing cyanobacteria hydrolysate as feedstock for the animal cells The cyanobacteria would be grown in an open pond made of concrete, harvested, sterilized, hydrolyzed and used as an animal cell growth medium. To these authors’ knowledge, this is not a technology or feedstock that is currently used for animal cell proliferation, nor is it one that is currently near feasibility given the current technical challenges of ACBM production. An amendment to the original study was later published that acknowledged technical challenges that the original study did not address (Tuomisto et al., 2014). While the published amendment also examined different scenarios with different feedstocks and bioreactor combinations, the authors acknowledged the high levels of uncertainty inherent to these untested approaches (Tuomisto et al., 2014).

An additional ACBM LCA that provided an increased level of detail was published in 2015 (Mattick et al., 2015). However, a close examination of the assumptions reveals some significant shortcomings of this study as well (Zimberoff, 2022). The process assessed in the study assumes the use of soy protein hydrolysate as an amino acid source, neglects to apply specific consumption rates to estimate the utilization of basal media and amino acids, and proposes the use of corn starch microcarriers for cell proliferation (Mattick et al., 2015). Once again, these assumptions are not accurate representations of current ACBM production.

In sum, the existing LCA literature on ACBM does not provide reliable estimates of the environmental impact of current or near-term ACBM production. This study seeks to address this gap in knowledge and provide a meaningful understanding of the environmental consequences of ACBM production. The assessment is based on a detailed model of ACBM production that is entirely based on peer-reviewed TEAs of ACBM systems as well as an existing LCA of the most representative ACBM media currently in use (Humbird, 2021; Risner et al., 2020). Given the existing level of investment, technological forecasting, and public funding associated with ACBM enterprises, this type of detailed environmental assessment of near-term ACBM production is critically needed (Zimberoff, 2022).

## Methods

This LCA was conducted utilizing the ISO 14040 and 14044 standards (International Organization for Standardization, 2006b, 2006a). The work builds on existing process models developed in peer-reviewed TEAs of ACBM (Humbird, 2021; Risner et al., 2020) as well as an existing LCA of an animal cell growth medium (Risner et al., 2023). The ISO process requires the following steps for a complete LCA: identifying the goal and scope, conducting a life cycle inventory (LCI), calculating the life cycle impact assessment (LCIA) and ongoing interpretation of all components throughout the process.

### Goal and scope

An updated environmental assessment is needed given the high levels of uncertainty of previously conducted ACBM LCAs. We aim to identify environmental challenges that should be addressed before seeking to industrialize a new meat production technology with assumed environmental benefits. In accordance with the ISO 14040 and 14044 standards, we have chosen the functional unit of a single kilogram of ACBM (wet basis) to allow for comparison with a similar conventionally produced ground beef product and ACBM products produced utilizing different growth mediums.

### Lifecycle inventory assessment

The development of a process model is an important element in identifying the inputs and outputs of a system. The Risner et al. and Humbird TEAs are the most complete studies that contribute to our understanding of the ACBM production process at this time. This study leverages the best components of both TEA models to create the ACBM process model for our LCA. Both the Risner et al. and the Humbird TEAs highlight the importance of the growth medium in influencing the economic viability of future ACBM products. A study examining the environmental impact of animal cell growth media has been utilized to further enhance the quality of our environmental assessment of near-term ACBM (Risner et al., 2023).

#### Risner et al. TEA

The Risner et al. TEA estimated the required volume of growth medium based on cellular glucose consumption rates and did not examine cellular amino acid consumption rates at the time. However, animal cells must have an amino acid source. The theoretical limit of the mass balance of the amino acids provided and protein produced is 1:1. In reality, it is lower since amino acids are also used as an energy source as well as for nucleic acid production. Table 1.0 provides a breakdown of animal cell composition (Humbird, 2021).

**Table 1.0.** Animal cell composition

|  |  |
| --- | --- |
| Moisture content | 70% |
| Dry matter | 30% |
| Protein (dry basis) | 70% |
| Lipid (dry basis) | 15% |
| Carbohydrate (dry basis) | 10% |
| Nucleic acid (dry basis) | 5% |

These are key assumptions for the new model which explores utilizing both the minimum glucose and amino acid requirements to generate minimum viability scenarios for our production system. We have taken the approach of utilizing a fed-batch system that supplies the cells with the nutrients in E8 as necessary. This approach allows for a concentrated feed to be added to bioreactor and prevents cells from experiencing issues related to osmotic pressures from increased nutrient concentrations. Risner et al. scenario 1 utilizes a glucose requirement would require 1,148 liters of E8 to produce a kilogram of ACBM. When E8 provision is scaled to match the amino acid requirements for cell cultivation, then it would require ∼292 liters of E8 to produce a kilogram of ACBM. Applying this amino acid requirement assumption to the previous UC Davis model shows that the Scenario 4 minimal requirement of E8 is actually not technically feasible, and Scenarios 2 and 3 may or may not be feasible depending on the protein content of the original inoculum. However, this limitation is accounted for in the updated model presented in this paper.

#### Humbird Technoeconomic assessment

To determine the cellular metabolic requirements, a “wild type” cellular metabolism and an “enhanced” cellular metabolism were examined. The wild type metabolism was deemed too inefficient for economic production due to lactate and ammonia production which inhibit cell growth. We only examined the enhanced cellular metabolism due to the economic concerns of the Humbird TEA. Equations 1 and 2 were utilized in the Humbird TEA to determine the mass of glucose, oxygen and amino acids needed for cellular proliferation. Dry cell matter (DCM) was determined, and the mass of each compound needed to produce a kg of ACBM (wet basis) was calculated.

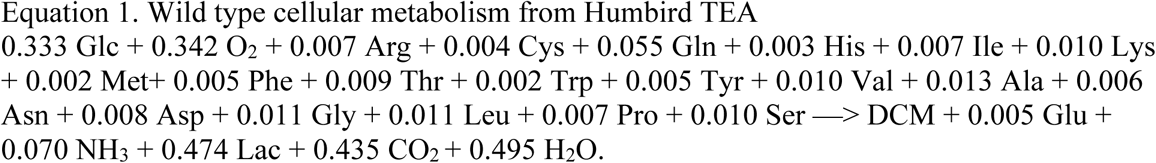

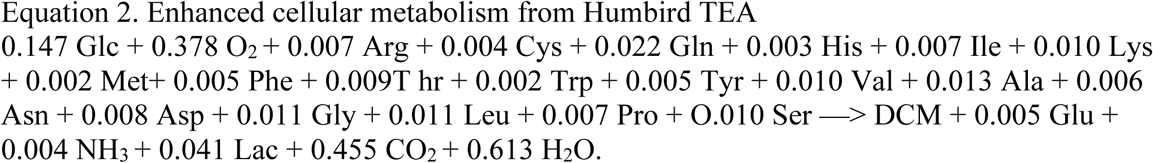

Both of Humbird’s cellular metabolism models assume the use of glutamine, but glutamine is not an E8 component. To address this gap, a literature source was used to determine microbial yield (0.368 g/g glucose) and the microbial method described in the E8 LCA was applied to determine the environmental impact of glutamine inclusion in the growth medium (Lv et al., 2021). It is likely not included in E8 due to stability issues; however, it plays an important role in cellular metabolism (Lu et al., 2019). Masses of minor protein ingredients such as insulin, transferrin, fibroblast growth factor (FGF) and transforming growth factor (TGF) were also accounted for on a functional unit basis.

The Humbird TEA also accounted for the power usage per batch. We examined the energy usage based upon batches per year (54,000 batches per year at 1852 kg/batch). Table 1 provides energy usage and unit conversions. This was then examined on a functional unit basis of 1 kg of ACBM.

**Table 1.**
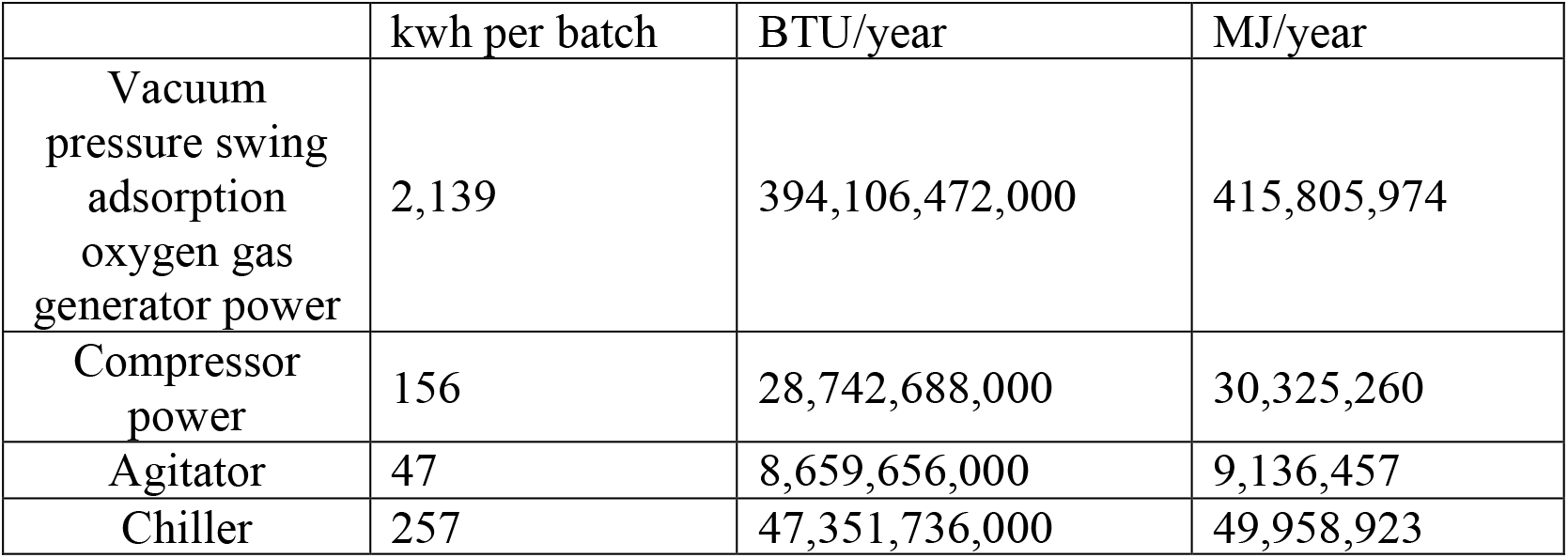
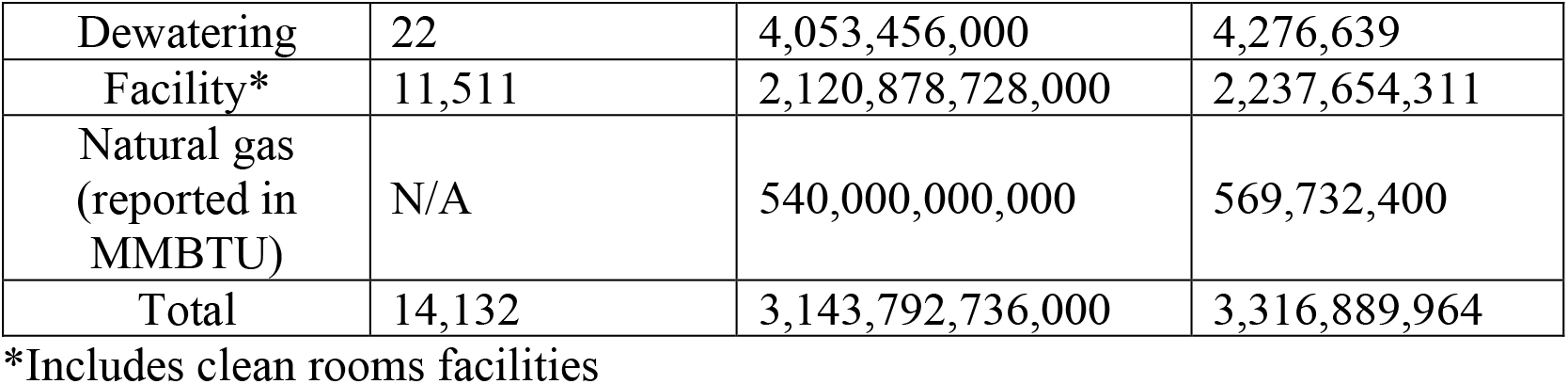
Humbird TEA energy estimates

In sum, it was determined that the Humbird TEA had more complete accounting of energy use and capital expenditures than the Risner et al. TEA, but Humbird TEA assumptions about the growth medium needed to be updated to include additional necessary vitamins and minerals for animal cell growth.

#### Combined production system

Utilizing the best information from the Risner et al. TEA, the Humbird TEA, and the E8 LCA a new production system was modeled to understand the near-term impact of ACBM production. The capital expenditures described in the Humbird TEA were complete, however these were not considered for this assessment e.g. mining of metals for bioreactor construction. The energy requirements from the production facility modeled from Humbird were used to estimate potential energy requirements. Figure 1 is a process flow diagram of a fed-batch ACBM production system with associated energy requirements.

**Figure 1.**
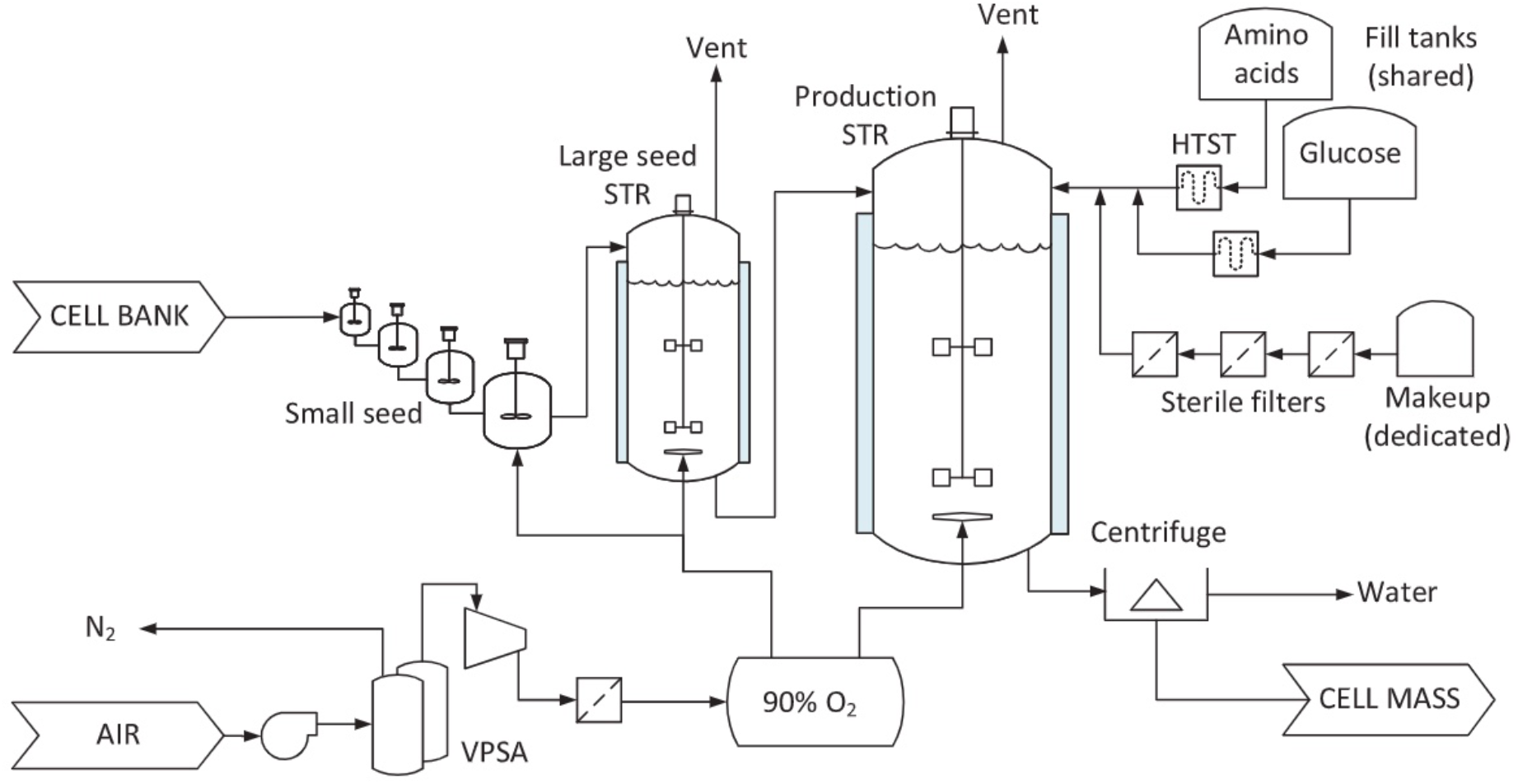
Fed-batch ACBM production system utilized in this LCA of ACBM This image was taken from Scale-up economics for cultured meat (Humbird, 2021)

The use of glucose consumption rate and required amino acid content was taken from the Risner et al. TEA. Also, the idea of utilizing a highly refined growth medium can be attributed to the Risner et al. TEA. The Humbird growth medium requirements were also calculated and utilized for scenario development. The growth medium requirements were entered into OpenLCA which contained the datasets for the E8 growth medium components. Figure 2 provides a map of how the main literature sources were utilized to provide an updated LCA of ACBM.

**Figure 2.**
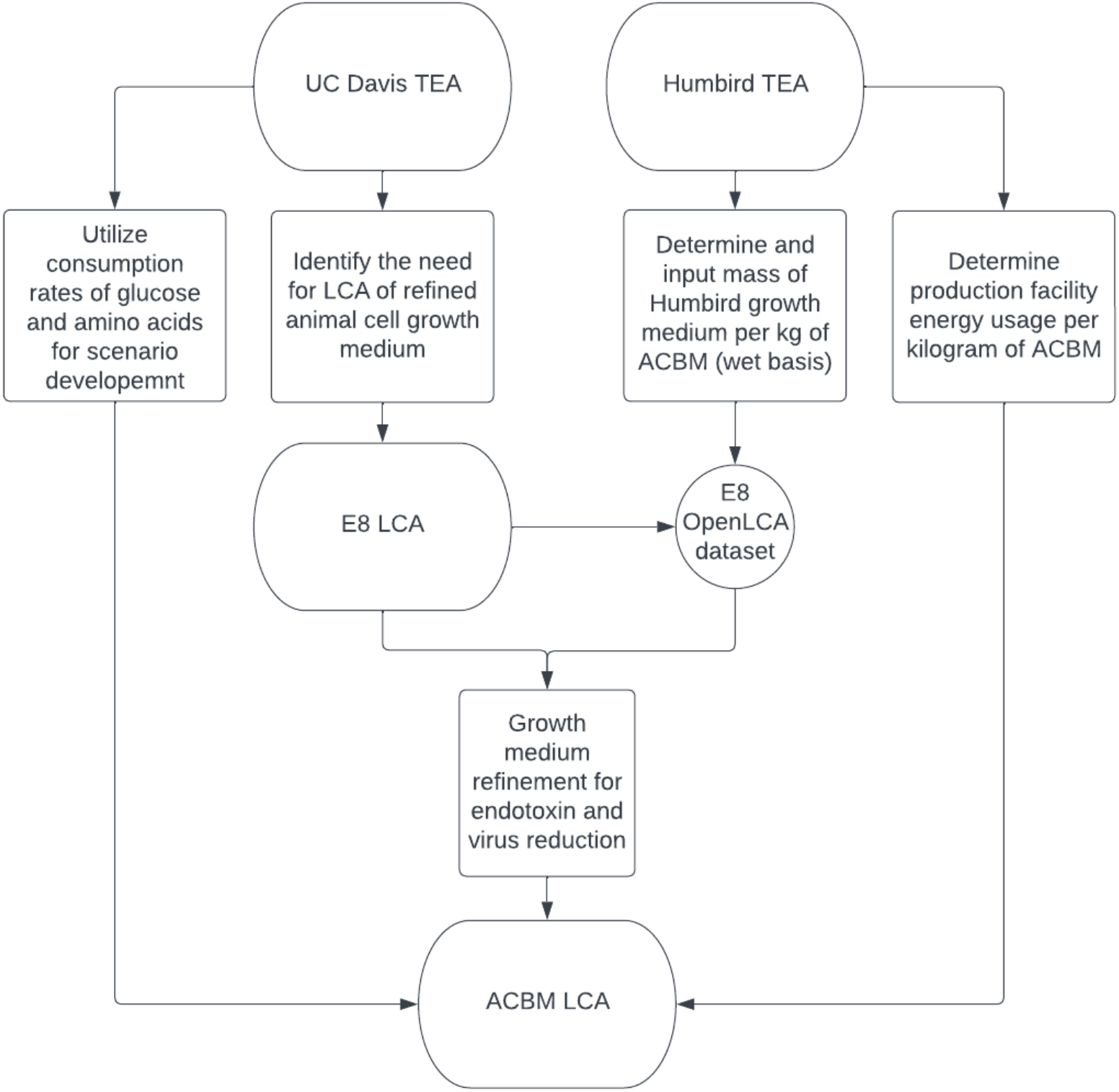
ACBM LCA main literature source map

#### Lifecycle Impact assessment (LCIA)

After all the inputs were identified and consolidated, a life cycle impact assessment was completed utilizing data and methods from the E8 LCA, OpenLCA v.1.10 software and OpenLCA LCIA v2.1.2 methods software. The tool for reduction and assessment of chemicals and other environmental impacts (TRACI) 2.1 was the LCIA methods utilized in the OpenLCA LCIA software, and these results were combined with the facility power data to determine the potential environmental impact of the production of 1 kg ACBM (wet basis).

#### Scenario analysis

All scenarios utilize a fed-batch system as described in the Humbird (2021) TEA. Energy estimates from the Humbird TEA are utilized in all scenarios. Growth medium components were assumed to be delivered to the animal cells as needed and the build-up of growth inhibiting metabolites such as lactate or ammonia are not accounted for unless specifically stated in the scenario. The growth medium substrates are also assumed to be supplied via fed batch to achieve the highest possible specific growth rate in the production bioreactor. The three minimum/base scenarios were defined utilizing data from the Risner et al. and Humbird TEAs then a purification factor was applied based on the results from a LCA which examined the environmental impact of fine chemical and pharmaceutical production (Wernet et al., 2010).

Each of the three base scenarios were examined independently and then with the purification factor applied for a total of six scenarios in the assessment (see descriptions below):

- **Risner et al. glucose consumption rate (GCR) scenario**: Reported estimates of the cellular glucose consumption rate were utilized to estimate the required growth medium volume in the Risner et al. TEA. This is same nutrient requirement as Scenario 1 from the Risner et al. TEA, however it is being delivered in a fed-batch manner as described by the Humbird system. The entire volume of growth medium is not assumed to be replaced, but the required nutrients are added as needed. This scenario utilizes E8 for its growth medium and it is estimated to require the equivalent of 1,148 L of E8 to produce one kilogram of ACBM wet basis.
- **Risner et al. amino acid requirement (AAR) scenario**: This scenario utilizes E8 as its growth medium and provides the minimum amount of amino acids needed to achieve the minimum amount of cellular protein mass for one kilogram of ACBM to be produced. This scenario indicates that 291.5 liters of E8 would contain the necessary amount of amino acids to produce a kilogram of ACBM wet basis with 21% (w/w) protein content.
- **Humbird growth medium scenario (HGM):** This scenario utilizes the Humbird TEA enhanced metabolism equation (equation 2) to estimate the total required growth medium nutrients. The wild-type metabolism was not utilized for scenario development due to it being deemed economically unfavorable. This scenario utilizes 0.35 kg of glucose, 0.16 kg of oxygen, 0.26 kg of amino acids, and minor protein ingredients (209.52 mg of insulin, 115.56 mg of transferrin, 1.08 mg of FGF and 0.02 mg of TGF) to produce one kg of ACBM wet basis.
- **Purification factor (PF):** Fine chemical or pharmaceutical production is more energy and resource intensive than bulk chemical production (Wernet et al., 2010). To account for this, the authors have utilized an LCA which compared fine chemical production to bulk chemical production. It was reported the cumulative energy demand (MJ) was 20x greater than bulk chemical production and the global warming potential (GWP) was 25x greater than bulk chemical production (Wernet et al., 2010). In animal cell culture, growth mediums are highly refined to prevent contamination from endotoxin and other contaminates. Given the resource intensity of fine chemical production, a purification factor of 20x is utilized to account for the resources associated with high levels of refinement.

The scenarios were developed to examine a range of potential environmental impacts utilizing the information available to the authors. As more complete information is ascertained about

ACBM production, additional scenarios could be developed to provide a more complete understanding of the true environmental impact of the ACBM products.

## Results

The LCIA was conducted on both the base scenarios and scenarios with purified growth medium components. TRACI 2.1 results are shown in Table 2. The GWP for all ACBM scenarios (19.2 to 1,508 kg of CO_2_e per kilogram of ACBM) was greater than the minimum reported GWP for retail beef (9.6 kg of CO_2_e per kg of FBFMO) (Poore & Nemecek, 2018). The GWP of all purified scenarios ranged from 246 to 1,508 kg of CO_2_e per kilogram of ACBM which is 4 to 25 times greater than the median GWP of retail beef (∼60 kg CO_2_e per kg of FFBMO). Without purification of the growth medium components, the GWP of the GCR scenario is approximately 25% greater than reported median of GWP of retail beef (Poore & Nemecek, 2018).

**Table 2.** TRACI 2.1 LCIA results for each unprocessed and purified growth medium scenarios

|  | GCR | GCR-PF | AAR | AAR-PF | HGM | HGM-PF |
| --- | --- | --- | --- | --- | --- | --- |
| Smog (kg O<sub>3</sub> eq) | 4.5 | 89.4 | 1.1 | 22.7 | 0.69 | 13.8 |
| Acidification (kg SO<sub>2</sub> eq) | 0.6 | 12.9 | 0.2 | 3.3 | 0.10 | 1.9 |
| Respiratory effects (kg PM<sub>2.5</sub> eq) | 0.1 | 1.6 | 0.0 | 0.4 | 0.01 | 0.3 |
| Non carcinogenic (CTUh) | 0.0 | 0.0 | 0.0 | 0.0 | 0.00 | 0.0 |
| Ecotoxicity (CTUe) | 1,848.9 | 36,977.9 | 469.6 | 9391.7 | 229.92 | 4,598.4 |
| Global Warming (kg CO<sub>2</sub> eq) | 75.4 | 1,508.3 | 19.2 | 383.1 | 12.31 | 246.1 |
| Ozone depletion (kg CFC-11 eq) | 0.0 | 0.0 | 0.0 | 0.0 | 0.00 | 0.0 |
| Carcinogenics (CTU) | 0.0 | 0.0 | 0.0 | 0.0 | 0.00 | 0.0 |
| Eutrophication (kg N eq) | 0.5 | 9.0 | 0.1 | 2.3 | 0.07 | 1.4 |
| Fossil Fuel depletion (MJ surplus)* | 85.3 | 1,706.4 | 21.7 | 433.4 | 14.89 | 297.8 |

\*Energy usage by ACBM production facility not accounted for in the table

It should be noted that the system boundary of this LCA stops at the ACBM production facility gate and does not include product losses, cold storage, transportation, and other environmental impacts associated with the retail sale of beef. Inclusion of these post-production processes would increase the GWP of ACBM products. Figure 3 illustrates the difference in the GWP of retail beef and cradle to upstream ACBM production gate.

**Figure 3.**
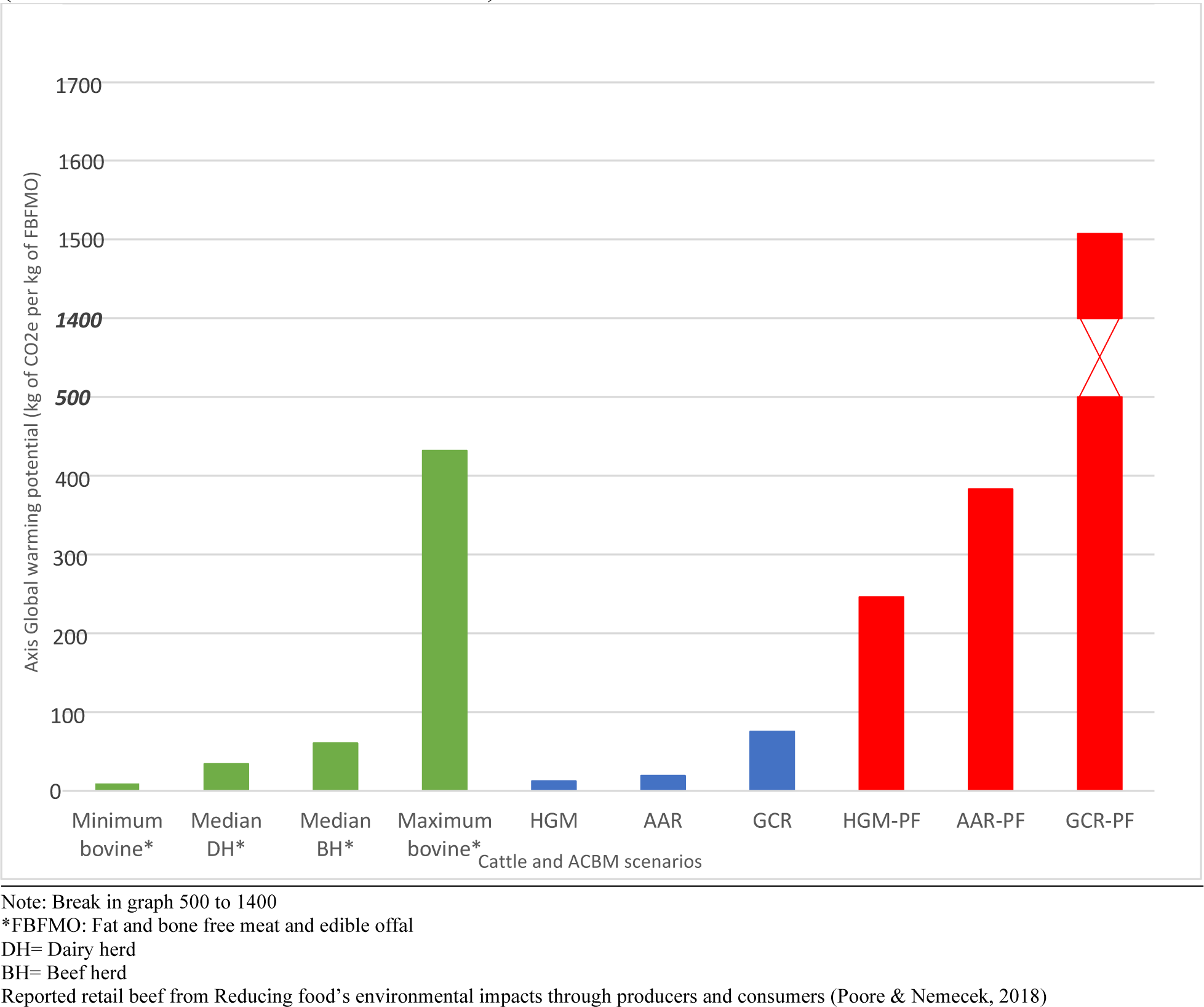
Comparison of GWP of the ACBM production scenarios and reported retail beef values (fat and bone free meat and edible offal).

The fossil fuel depletion metrics were greater for all the ACBM production scenarios as compared to the low boneless beef metric (see figure 4). For unpurified scenarios, the higher level of energy use is largely associated with upstream processing facilities producing input products required for ACBM production. The HGM scenario was approximately ∼1 MJ per kilogram greater than the lower estimate for boneless beef (Maysami & Berg, 2021). The AAR-PF and AGM-PF scenarios with growth mediums refined for animal cell culture required approximately an order of magnitude more energy than the reported low for boneless beef. The high cumulative energy demand for boneless beef was approximately double the fossil fuel depletion of the AAR and HGM scenarios. The fossil fuel depletion for scenarios with purified growth medium components were approximately 3 to 17 times greater than the reported high for boneless beef.

**Figure 4.**
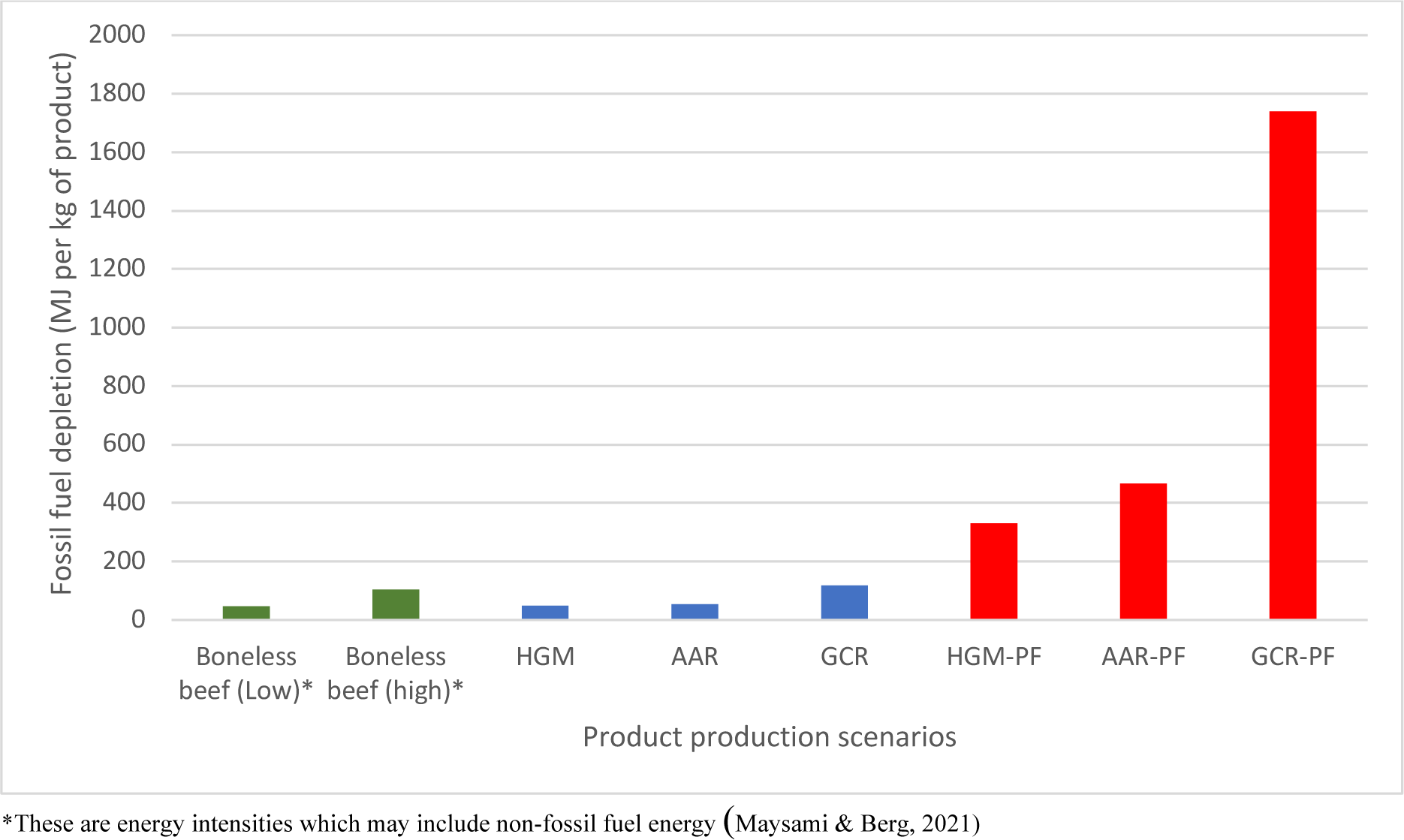
Fossil fuel depletion of each ACBM production scenario in comparison with boneless beef

Our system boundary for ACBM production does not include post-harvest handling, storage and transport which all require energy in some form. These additional energy inputs may increase the energy intensity/fossil fuel depletion of ACBM products indicating the reported results may be viewed as minimums.

## Discussion

Our results indicate that ACBM is likely to be more resource intensive than most meat production systems according to this analysis. In this evaluation, our primary focus has been on the resource intensity of the growth mediums. We have largely focused on the quantity of growth medium components (e.g. glucose, amino acids, vitamins, growth factors, salts, and minerals) and attempted to account for purification requirement of those components for animal cell culture. We also acknowledge that our analysis may be viewed as minimum environmental impacts due to several factors including incomplete datasets, the exclusion of energy and materials required to scale the ACBM industry and exclusion of the energy and materials needed to scale industries which would support ACBM production.

We examined the growth mediums utilized in both UC Davis and Humbird TEAs and selected the UC Davis TEA as a more reasonable assumption given its more complete composition.

Figure 5 compares the global warming potential of the different categories of basal growth medium components within each growth medium and illustrates differences in the basal mediums, such as the inclusion of vitamins, inorganic salts, and other components in the E8 growth medium.

**Figure 5.**
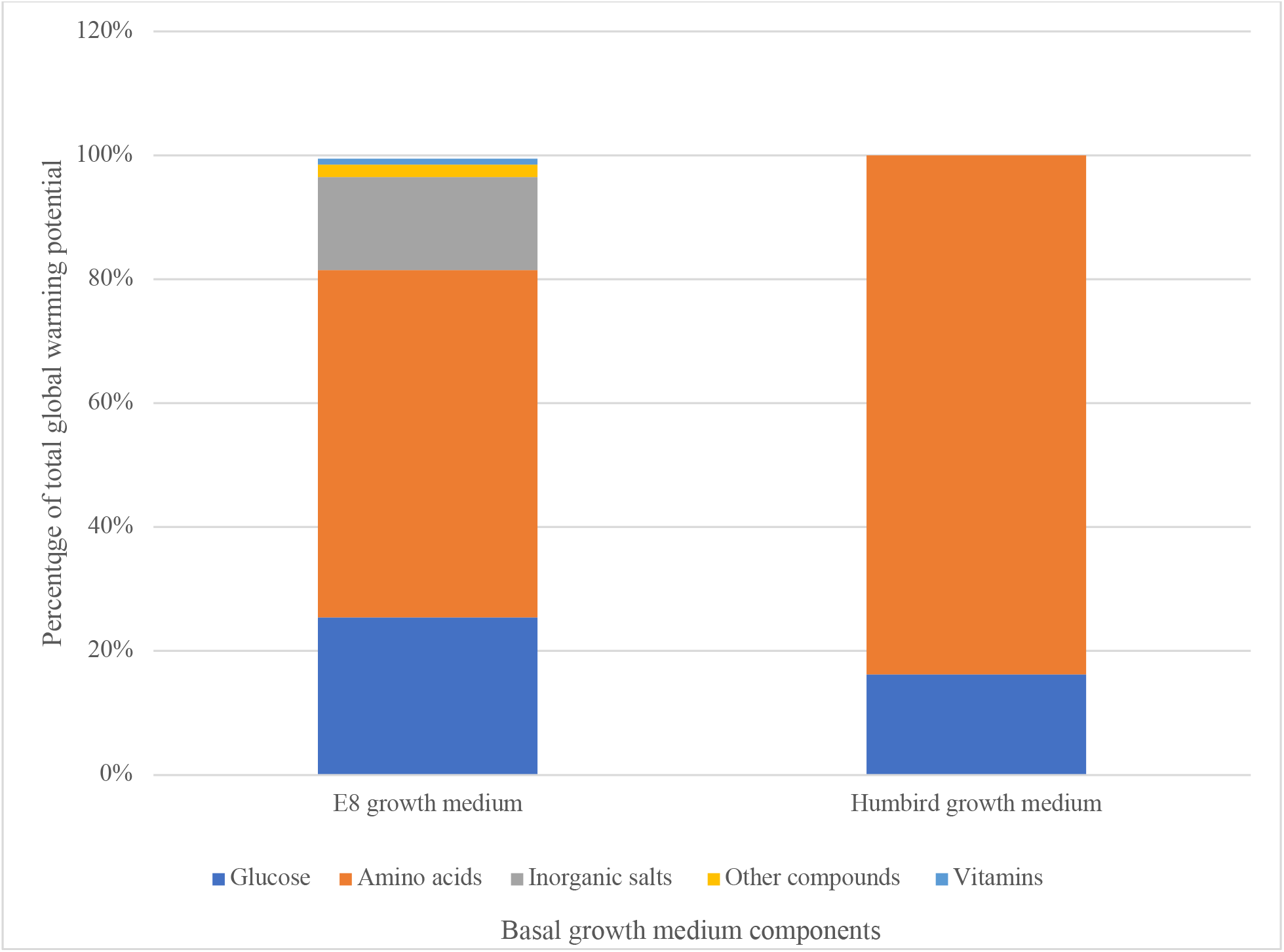
Growth medium component contribution to global warming potential of each basal growth medium

Given the stringent medium component purity requirements for animal cell culture, the high purification scenarios with E8 as the growth medium are likely to represent the more accurate environmental impact of ACBM production. It should also be noted that these results should be considered a minimum since the E8 LCA is admittedly non-exhaustive (Risner et al., 2023). The E8 LCA does not account for 4-(2-hydroxyethyl)-1-piperazineethanesulfonic acid (HEPES) and lipoic acid production and there is only partial accounting of the embedded resources and energy for other E8 components (Risner et al., 2023). Scenarios AAR and AAR-PF assume a 100% conversion of amino acids to protein. This assumption is probably an unrealistic assumption given the amino acids also supply the nitrogen atom and amino group in the synthesis of nucleotide bases and nitrogen-containing sugars (Hu, 2020). The amino acid carbon skeleton is also utilized in the formation of groups like the functional methyl group (Hu, 2020). This indicates that AAR-PF may be an unlikely minimum as well.

Animal cell culture is inherently different than culturing bacteria or yeast cells due to their enhanced sensitivity to environmental factors, chemical and microbial contamination. This can be illustrated by the industrial shift to single use bioreactors for monoclonal antibody production to reduce costs associated with contamination (Jacquemart et al., 2016). Animal cell growth mediums have historically utilized fetal bovine serum (FBS) which contains a variety of hormones and growth factors (Jochems et al., 2002). Serum is blood with the cells, platelets and clotting factors removed. Processing of FBS to be utilized for animal cell culture is an 18-step process that is resource intensive due to the level of refinement required for animal cell culture. Thus, the authors believe that commercial production of an ACBM product utilizing FBS or any other animal product to be highly unlikely given this high level of refinement.

The requirement of endotoxin removal would also contribute to the environmental impact of ACBM products which makes our LCIA results for the minimum scenarios to be underestimated minimums. Utilization of commodity grade growth medium components such as glucose for animal cell growth is unlikely unless the components undergo an endotoxin separation process. The effect of endotoxin can vary greatly depending on cell type and source; however 25 ng/ml of endotoxin was shown to cause cell apoptosis when coupled with non-lethal heat shock (Corning, 2020). Concerns over endotoxins may also limit the possibility of utilizing novel amino acid sources (e.g. plant hydrolysates) in the ACBM media since endotoxin is amphiphilic with its hydrophilic polysaccharide fraction and hydrophobic lipid fraction. The amino acids in the plant hydrolysate will interact differently depending upon their functional properties (e.g., hydrophobicity, charge). This multitude of interactions will potentially make separation difficult without additional processing steps, which will further increase the environmental impact of the ACBM growth medium and subsequently ACBM products. However, endotoxin does have an overall negative charge which may be beneficial for separation, For these reasons, the authors believe that scenarios which account for purification to be closer to a true minimum rather than the minimum baseline scenarios. An additional strategy for potential ACBM producers would be to develop cell lines which are endotoxin tolerate which may be help reduce the potential environmental impact of ACBM products.

We did not consider the environmental impact of scaling up ACBM production facilities. In 2021, the total cell culture bioprocessing capacity was 17.4 million liters with mammalian cell culture capacity being 11.75 million liters (Langer & Rader, 2021). The Humbird TEA states that each fed-batch production facility would require a total bioreactor volume of 649 m^3^ and that it would require ∼14.7 identical facilities to produce 100,000,000 kg of ACM annually, or an additional 9,540,300 liters of mammalian cell culture capacity. If this capital expansion was included in our LCA, we would need to expand our system boundary to include all the resources used in the mining of the materials and construction of these facilities. We also have not included the environmental impacts associated with scaling up multiple production facilities to produce the required mass of growth media components necessary for ACBM production at scale (Humbird, 2020, 2021). For these reasons, we believe that additional work is necessary to provide this expanded view of the environmental impact of producing ACBM at scale.

## Conclusion

Critical assessment of the environmental impact of emerging technologies is a relatively new concept, but it is highly important when changes to societal-level production systems are being proposed (Bergerson et al., 2020). Agricultural and food production systems are central to feeding a growing global population and the development of technology which enhances food production is important for societal progress. Evaluation of these potentially disruptive technologies from a systems-level perspective is essential for those seeking to transform our food system. Ideally, systems-level evaluations of proposed novel food technologies will allow policymakers to make informed decisions on the allocation of government capital. Proponents of ACBM have hailed it as an environmental solution that addresses many of the environmental impacts associated with traditional meat production. Upon examination of this highly engineered system, ACBM production appears to be resource intensive when examined from the cradle to production gate perspective for the scenarios and assumptions utilized in our analyses.

The existing LCAs of ACBM are insufficient for assessing the environmental impact of this emerging food technology. The main issue with these preeexisting studies is that their technology models do not accurately reflect the current/near term practices which will be utilized to produce these products. Our environmental assessment is grounded in the most detailed process systems available that represent current state-of-the-art in this emerging food technology sector. Our model generally contradicts these previous studies by suggesting that the environmental impact of cultured meat is likely to be higher than conventional beef systems, as opposed to more environmentally friendly. This is an important conclusion given that investment dollars have specifically been allocated to this sector with the thesis that this product will be more environmentally friendly than beef.

Given this assessment, investing in scaling this technology before solving key issues like developing an environmentally friendly method for endotoxin removal or adapting cell lines which are endotoxin resilient would be counter to the environmental goals which this sector has espoused. Perhaps a focus on advancing these precompetitive scientific advances might lead to a better outcome for all. For example, solving the endotoxin challenge would also substantially benefit the biomedical and biopharmaceutical industries and their consumers by substantially reducing the cost of production. Another example would be the development of a technological innovation that allows for the use of an inexpensive animal cell growth media produced from agricultural by-products. In short, our environmental assessment highlights the need for critical, detailed environmental assessments of emerging technologies to guide governmental agencies and the private sector *in advance of* allocating substantial research funding towards initiatives that assume transformational environmental benefits in the absence of rigorous analysis.

In sum, understanding the minimum environmental impact of near term ACBM is highly important for governments and businesses seeking to allocate capital that can generate both economic and environmental benefits (Zimberoff, 2022). We acknowledge that our findings would likely be the minimum environmental impact due to the preliminary nature of our LCA. This LCA aims to be as transparent as possible to allow the interested parties to understand our logic and why we have developed these conclusions. We also hope that our LCA will provide evidence of the need for additional critical environmental examination of new food and agriculture technologies.

